# Fitness cost associated with enhanced *EPSPS* gene copy number and glyphosate resistance in an *Amaranthus tuberculatus* population

**DOI:** 10.1101/2021.01.09.426028

**Authors:** Helen M. Cockerton, Shiv S. Kaundun, Lieselot Nguyen, Sarah Jane Hutchings, Richard P. Dale, Anushka Howell, Paul Neve

## Abstract

The evolution of resistance to pesticides in agricultural systems provides an opportunity to study the fitness costs and benefits of novel adaptive traits. Here, we studied a population of ***Amaranthus tuberculatus*** (common waterhemp), which has evolved resistance to glyphosate. Following the production of seed families with contrasting levels of glyphosate resistance, we assessed the growth and fitness of seed families in the absence of glyphosate and determined their ability to compete for resources under intra- and interspecific competition. Further investigation revealed a positive correlation between the level of resistance and gene copy number for the 5-enolpyruvylshikimate-3-phosphate synthase (EPSPS) glyphosate target, thus indicating gene amplification as the mechanism of resistance within the population. Resistant common waterhemp plants were found to have a lower competitive response when compared to the susceptible phenotypes. A substitution rate of 2.76 glyphosate resistant plants was required to have an equal competitive effect as a single susceptible plant. A growth trade-off was associated with the gene amplification mechanism under intra-phenotypic competition where 20 extra gene copies were associated with a 26.5 % reduction in dry biomass. Interestingly, this growth trade-off was mitigated when assessed under interspecific competition from maize.

## Introduction

Crop protection practices are designed to limit the incidence and impacts of crop pests, including weeds. Unfortunately, these practices also provide a selection pressure for pest organisms to overcome control strategies. Pathogens, pests and weeds have all evolved resistance to pesticides through selection of ***de novo*** mutations and/or standing genetic variation (Hawkins ***et al.*** 2018). Evolutionary theory states that adaptation to resist or tolerate a novel stress may incur a fitness cost when the individual is returned to the original “stress-free” environment (Herms & Mattson 1992; Bergelson & Purrington 1996; Coustau, Chevillon & ffrench-Constant 2000; Fry 2003; Vila-Aiub, Neve & Roux 2011; Vila-Aiub, Gundel & Preston 2015). Ultimately, the quantification of fitness costs in a herbicide resistant population can inform resistance management strategies in order to assist effective weed control (Cousens & Mortimer 1995; Jordan 1999; Vila-Aiub, Neve & Powles 2005a).

The size of the fitness benefit determines the rate at which a resistance allele will establish within a population under selection (Roux ***et al.*** 2006). Similarly, the presence of a resistance cost will determine if the frequency of a resistance allele will reduce in the absence of selection (Coustau ***et al.*** 2000). Therefore, the benefit-cost fitness balance is thought to determine the frequency of a resistance allele under different agricultural management regimes (Roux ***et al.*** 2006; Vila-Aiub ***et al.*** 2011).

The fitness cost associated with an adaptive allele may manifest as a direct cost through a pleiotropic effect of the causative allele itself, or via ecological trade-offs in life history traits such as biomass (McCloskey & Holt 1990), emergence date (Vila-Aiub ***et al.*** 2005b), height and flowering time (Roux, Matejicek & Reboud 2005) each of which may ultimately result in a direct fitness cost within a resource limited environment (Délye, Jasieniuk & Le Corre 2013). Fitness costs have been widely reported for antibiotic resistance (Lee & Edlin 1985; Nguyen, Phan & Lp 1989; Bentley ***et al.*** 1990; Purrington & Bergelson 1997). However, such costs are not universally observed for herbicide resistant weed populations, and where costs have been observed they have been shown to be influenced by the genetic background of the population (Paris ***et al.*** 2008; Vila-Aiub, Neve & Powles 2009). It is believed that fitness costs, if present, can be magnified by intra and inter-specific competition for resources (Bergelson & Purrington 1996).

The mechanism of evolved resistance may also influence the magnitude and type of fitness cost that is expressed (Paris ***et al.*** 2008). For example, resource allocation theory states that a resistant individual which metabolises xenobiotics will divert resources away from growth and reproduction and towards defence, resulting in a cost of adaptation (Bergelson & Purrington 1996). Furthermore, additional gene copy numbers are associated costs due to allocation of resources towards over-production of an amplified protein (Tang & Amon, 2012). By contrast, a mutation in a pesticide target enzyme may impact on the efficacy of enzyme function (Vila-Aiub ***et al.*** 2005b; Tardif, Rajcan & Costea 2006; Menchari ***et al.*** 2007), leading to fitness costs through the disruption of normal metabolic processes.

***Amaranthus tuberculatus*** (Common Waterhemp) is a prevalent, problematic weed in the USA and Canada. Over the last three decades genetically-modified glyphosate tolerant crops have been grown extensively, resulting in multiple annual applications of glyphosate. Indeed, the major selling point of glyphosate tolerant crops is the provision of a low maintenance one-step weed management strategy. A prolonged reliance on glyphosate tolerant crops has provided the strong selection pressure required for the evolution of glyphosate resistant weed populations (Neve, 2008). By 1998, variable glyphosate responses had been identified in ***A. tuberculatus*** field populations (Zelaya & Owen, 2002, 2005), but it was ten years before the first confirmed incidence of field-evolved glyphosate resistant populations (Legleiter & Bradley 2008). Today, glyphosate resistant ***A. tuberculatus*** has been documented in 21 states of the USA and in Ontario, Canada (Heap, 2020). Canadian populations have established from multiple origins, both through gene flow from bordering epidemics and through de-novo evolution of resistance (Kreiner ***et al.*** 2019).

Here we studied a glyphosate resistant population of common waterhemp to establish the mechanism of glyphosate resistance and the presence of fitness costs associated with that mechanism. We test the hypothesis that resistance is associated with a fitness cost in the absence of glyphosate by carrying out two competition experiments. A response surface experiment was used to compare the competitive response and effect of resistant and susceptible phenotypes. Whereas a neighbourhood design experiment was used to determine if there was a trade-off between seed family level resistance and resistance cost under intra and inter specific competition with maize.

## Materials and Methods

### Plant material

Common waterhemp seed was collected from glyphosate-resistant soybean fields (N 44.78, W 95.21) in Renville, Minnesota, USA. Original seed collection was performed in 2007 after four glyphosate applications resulted in poor weed control. The generation of resistant and susceptible seed families from a single population permits fitness parameters to be studied through limiting background genetic variation (Vila-Auib et al., 2011). Resistant seed families were generated through crossing 10 resistant female common waterhemp plants with 10 resistant male common waterhemp plants in a bulk cross, susceptible seed families were generated in a similar fashion. Parental common waterhemp plant resistance status was identified through phenotyping of vegetative clones as outlined in Figure 1. After cross pollination, seed extracted from a single plant formed a seed family. A subsequent dose response experiment confirmed resistance levels of seed families and six families were selected to represent a range of glyphosate resistance levels amongst seed families.

**Figure 1:**
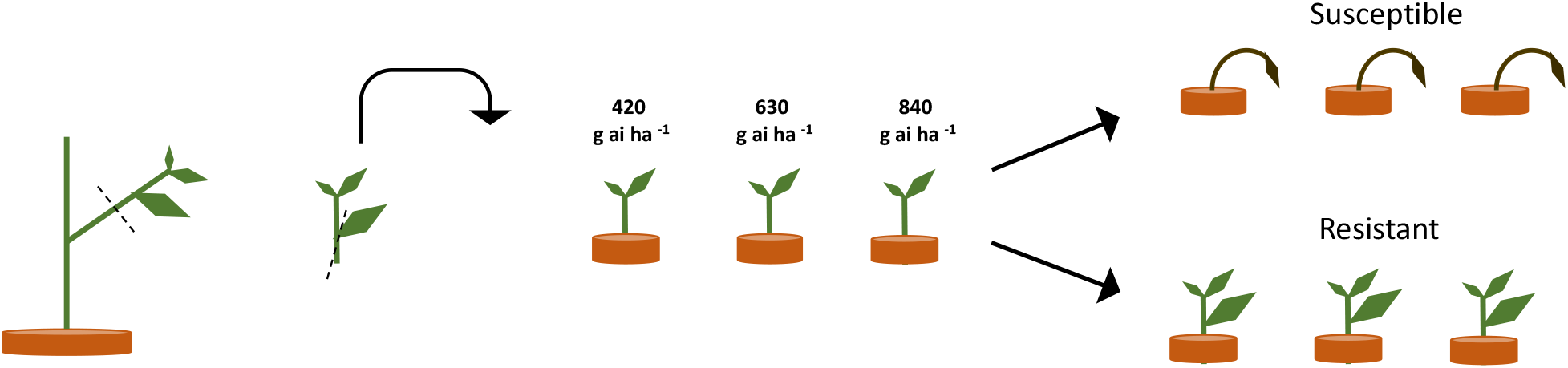
Scheme of clone propagation method for parental plant herbicide phenotyping. Three leaf cuttings were removed from individual parental plants and transplanted into pots to encourage rooting and re-growth. 18 days after propagation, three established clones per plant were treated with 420, 640 or 840 g ai ha^−1^ of glyphosate. Survival and growth were assessed and plants were scored as resistant or susceptible depending on plant survival, mortality and glyphosate symptoms at the three glyphosate doses.

### Seed family dose response resistance quantification

***A. tuberculatus*** seed from 6 glyphosate-selected seed families (above) was imbibed on Levington growing media (FS2) at 5 ^o^C for 7 d. and germinated in a glasshouse at 24:18 °C, 16:8 hr (day: night), with supplementary lighting. Seedlings were transplanted 14 days post germination into 1.5 L pots (10 per pot) containing medium grade sphagnum moss peat and pots were maintained in the glasshouse. Six seed families were assessed across three replicate pots per treatment, with a total of 360 plants in 36 pots per block arranged in a randomized block design. The S2 population was used as a sensitive control, S2 was sourced from Azlin Seed Service and had a similar morphology to the resistant populations. Glyphosate (isopropylamine glyphosate salt) was applied at the 6-8 leaf stage with 0, 105, 210, 420, 840 or 1680 g ai ha^−1^ using a Berthoud Velmorel 200 pro knapsack sprayer and Deflector Anvil Polijet nozzle (D/1.2/1) at 200 kPa, 2 km hr^−1^ at 40 cm above the plant canopy to deliver 300 L ha^−1^. Percentage survival and above ground biomass was assessed 21 days after treatment. Mortality was assigned based on observation of complete necrosis, apical meristem necrosis, and root system disintegration. The lethal dose required to kill 50% of individuals (LD_50_) was determined for each seed family based on the best fitting dose response model (log logistic, Weibull 1 or Weibull 2) where goodness-of fit was determined using a Pearsons’s chi squared test. Analysis was conducted in R (R version 2.15.1, R Development Core Team, 2009) using the drc package (Ritz & Streibig, 2005). Resistance indices were calculated as the LD_50_ of the putative resistant population as a ratio of the standard sensitive LD_50_ (LD_50_R / LD_50_S).

### Resistance mechanism determination

Elucidating the mechanism of resistance present in a population provides genotypic context for phenotypic observations. Target site sequencing and impaired translocation were assessed as detailed in the supplementary methods. Gene copy number (GCN) quantification was assessed through qPCR as detailed below.

DNA extraction was achieved through grinding approximately 0.5 g of common waterhemp leaf material in a pestle and mortar with 3 ml of grinding buffer (100 mM NaOAc pH 4.8; 50 mM EDTA pH8; 500 mM NaCl; 2% PVP; 1.4% SDS; H_2_O) and incubated at 65 °C for 15 min, 1 ml of ammonium acetate was added to supernatant and incubated at 65 °C for 10 min. Polysaccharide contaminants were removed by two subsequent additions of phenol: chloroform: iso-amyl alcohol (25:24:1) (pH 8) and one chloroform: iso-amyl alcohol (24:1). After each addition, solutions were mixed by inversion and centrifuged at 13000 rpm for 5 min and the aqueous layer was kept. Precipitation of DNA was achieved through addition of 0.6 V of cold isopropanol and 0.1 V of 3M NaOAc, chilled at −20 °C for 30 min and centrifuged at 13000 rpm for 10 min. Supernatant was discarded, pellets were dried for 10 min and DNA was resuspended in TE.

5-enolpyruvylshikimate-3-phosphate synthase (***EPSPS***) gene copy number was determined through qPCR on genomic DNA. Master mixes of Takyon Master Mix for SYBR^®^ Assay mix (Eurogentec): Water: Forward primer: Reverse primer: DNA template (5:1:1:2:1.25). Primer sequences in Supp. Table 2 for the gene of interest ***EPSPS*** and housekeeping genes carbamoyl phosphate synthetase ***CPS*** & acetolactate synthase (***ALS***). Reactions were run in triplicate for 6-8 individuals per seed family on an Applied Biosystems^®^ 7500 Fast Real-Time PCR System. qPCR cycle conditions were 95 °C for 3 min, 40 cycles of 95 °C for 3 sec, 60 °C for 40 sec. Gene copy number (GCN) was calculated by Eq. 1 &2.

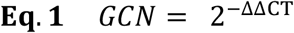

Where CT is the cycle threshold of ***EPSPS*** (E) or a combination of the control genes ***CPS*** and ***ALS*** (CA) and *ȳ* represents the standard sensitive geometric mean and 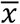 represents the seed family geometric mean. A regression analysis was conducted to determine the relationship between the average seed family GCN and whole plant glyphosate LD_50_.

### Response surface competition experiment

Plants from a resistant seed family (4.8 RI_LD50_) and susceptible seed family (1.7 RI_LD50_) were transplanted in a triangular grid at five densities of 1100, 560, 300, 200, 100 plants m^−2^ across seven resistant: susceptible proportions 6:0, 5:1, 4:2, 3:3, 2:4, 1:5, 0:6 (Figure 2). Rather than conducting a full factorial design representing all proportions and density combinations, 19 treatments were selected for study (Table 1). Three replicates boxes were produced per treatment (six replicates for 100 plants m^−2^) giving a total of 66 boxes arranged in a randomised block design. Plants were transplanted into boxes containing 13 kg of a 2:1 mix of top soil: Medium grade sphagnum moss peat (pH =7.6, K= 176.3, P= 80.4, NO3= 146.6, mg = 377.9 μg g^−1^). Border plants were included to eliminate edge effects, but were excluded from the analysis.

**Figure 2:**
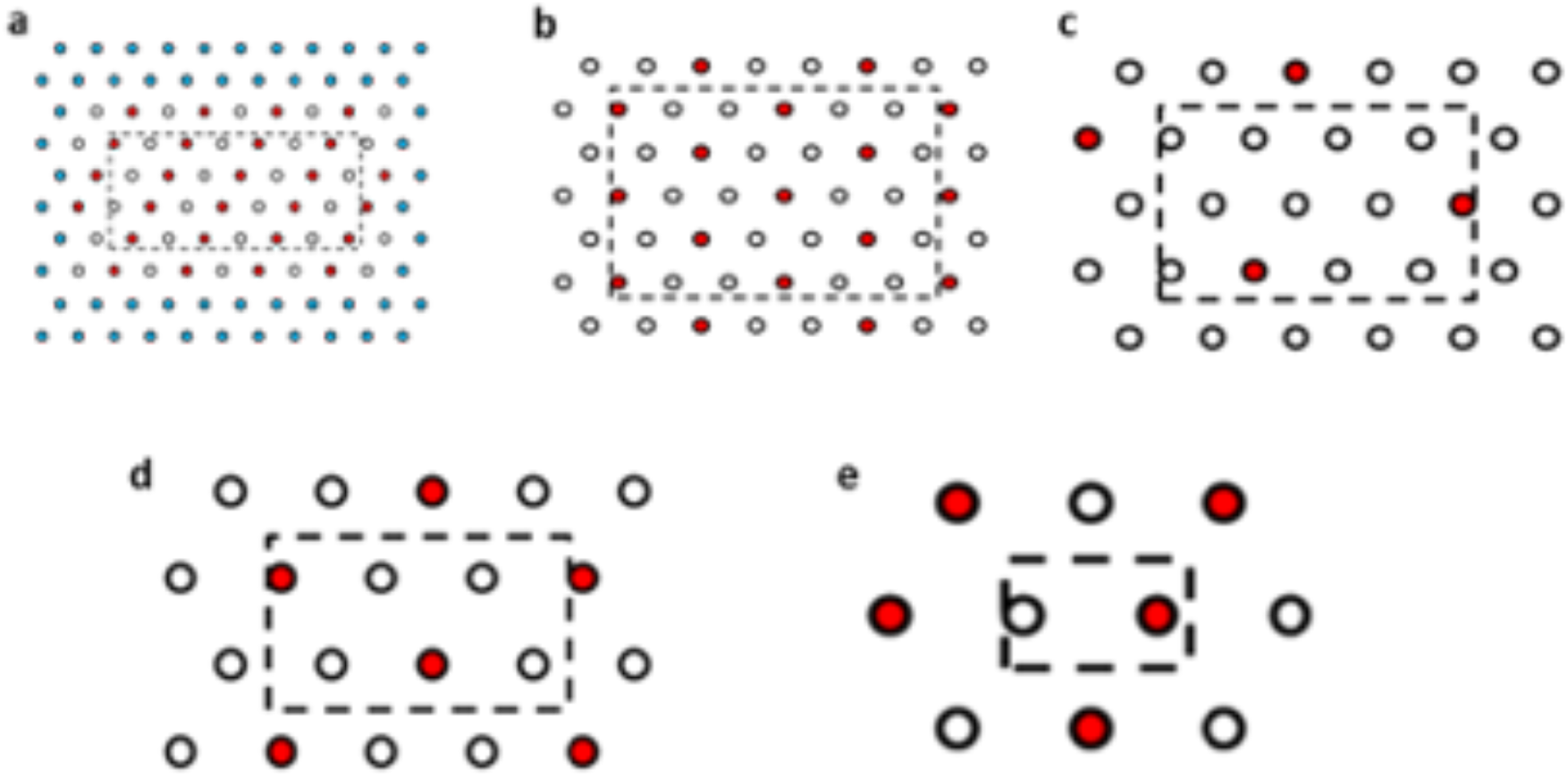
Planting arrangements at selected densities and proportions of resistant and susceptible seed families. a) R:S, 3:3 plant ratio, 1100 plants m^−2^, b) R:S 4:2, 560 plants m^−2^, c) R:S 6:1, 300 plants m^−2^, d) R:S 4:2 200 plants m^−2^, e) R:S 3:3 100 plants m^−2^. Red and white spots represent the plant position for plants from the most resistant and the most susceptible seed family phenotypes are interchangeable. Due to the high seedling volume required for the 1100 plants m^−2^ treatment, two rows of spacer plants (blue) were added to the outside of the layout, spacer plants were from seed family 153. Plants within the dashed line box were harvested, plants outside of the box acted as edge effect plants.

**Table 1:** Plant densities and resistant: susceptible (R:S) proportions used for the response surface experiment. + marks selected conditions.

| R:S | Density (plants m<sup>-2</sup>) |  |  |  |  |
| --- | --- | --- | --- | --- | --- |
|  | 1100 | 560 | 300 | 200 | 100 |
| 6:0 | + | + | + | + | + |
| 5:1 |  |  | + |  |  |
| 4:2 |  | + |  | + |  |
| 3:3 | + |  | + |  | + |
| 2:4 |  | + |  | + |  |
| 1:5 |  |  | + |  |  |
| 0:6 | + | + | + | + | + |

Above ground fresh biomass, sex, height and stem diameter were recorded at 114 days after transplanting. Log transformed biomass data was analysed in R (R version 2.15.1: 2012-06-22) (R Development Core Team, 2009) using the nonlinear least squares (nls) function in the stats package version 3.0.1. A hyperbolic curve (Pedersen et al., 2007, Equation 1) described the relationship between plant biomass, density and proportion and allowed the calculation of a, b and c parameters, where a is maximum plant biomass in the absence of competition, b is the competitive response and c is the relative competitive effect of phenotypes.

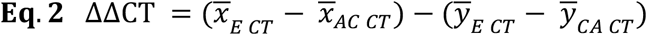

Effective density was calculated as ***N***_***i***_ + ***c***_***ij***_***N***_***j***_ where ***N*** is the density of the i or j phenotype and ***c***_***ij***_ is the ***c*** parameter determined by the either the ***i*** or ***j*** specific parameter. Parameters were compared using z-tests. Parameter c was tested for a difference from 1 using a t-test. Analysis included plants that have died as a result of competition.

### Neighborhood design competition experiment

Maize (cv. Kangaroo) and ***A. tuberculatus*** seedlings were transplanted as indicated in Figure 3, into 4 L pots containing 2 kg of the growth medium outlined above. Weed seedlings were transplanted so that cotyledons were 2 cm above the soil surface at the V1 corn growth stage. The three treatments denoted in Figure 3 allowed the investigation of the impact of ***A. tuberculatus*** on maize (***a*** & ***c***) and the impact of maize on ***A. tuberculatus*** (***a*** & ***b***). Layouts ***a*** & ***b*** are produced for the six ***A. tuberculatus*** seed families with contrasting levels of glyphosate resistance. In this way, layout ***a*** can determine the trade-off between plant growth and seed family resistance levels under interspecific competition and layout ***b*** can be used to assess this relationship under intraspecific competition. Replicate pots were arranged in a randomised block design, with ten replicates for all treatments, apart from maize-alone (***c***) which had 20 replicates. Competition experiments were conducted in a polytunnel from June to September. Total above ground and reproductive fresh biomass of maize and ***A. tuberculatus*** was recorded 107 days after transplanting. The overall impact of interspecies competition and biomass allocation was assessed through T-tests. Pearson correlations were calculated between biomass traits and resistant metrics, to determine trade-offs between fitness and resistance.

**Figure 3:**
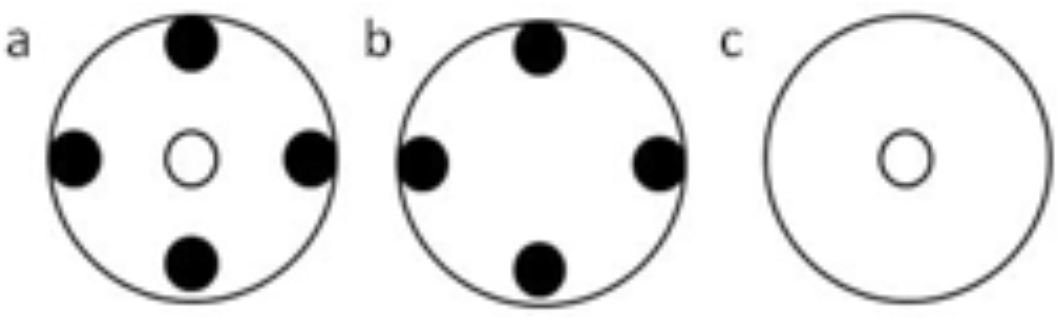
Layout of plants in neighbourhood design experiment. Large circles represent pots. Filled circles represent ***A. tuberculatus*** plants and open small circles represent maize plants.

## Results

### Quantitative segregation of glyphosate resistance

Dose response experiments were used to assess the herbicide resistant status of the common waterhemp field-collected population and the derived seed families. The field-collected population had a glyphosate resistance index of 3.2. Derived seed families had glyphosate resistance indices ranging from 1.7 to 5.3 (Table 2). The wide range of resistance phenotypes (levels of resistance) observed within and amongst seed families were suggestive of inheritance of a quantitative resistance trait. The variation in population-level resistance estimates provides a good range of seed families with which to test a trade-off between resistance level and fitness cost.

**Table 2.**
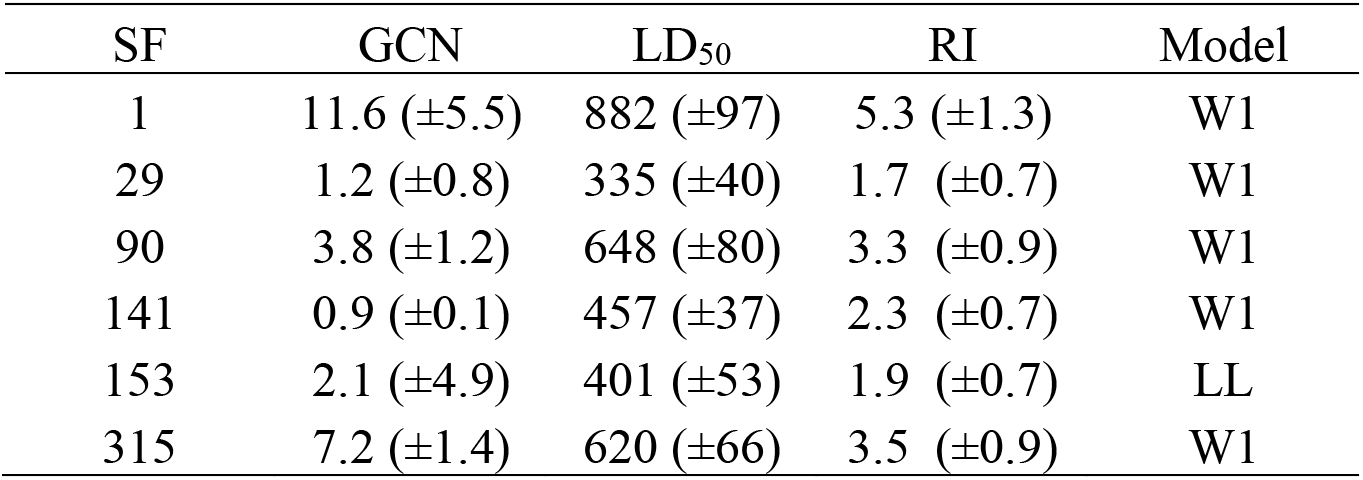
Seed family (SF) Gene copy number (GCN) of 5-enolpyruvylshikimate-3-phosphate synthase (***EPSPS***) relative to carbamoyl phosphate synthetase (***CPS***) and acetolactate synthase (***ALS***). Lethal dose required to kill 50 percent of individuals (LD_50_) Resistance Index (RI) relative to the field sensitive population S2. Standard errors are presented in parentheses. Model represents the best fitting dose response model; either log logistic (LL) or Weibull’s 1 (W1).

| SF | GCN | $LD_{50}$ | RI | Model |
| --- | --- | --- | --- | --- |
| 1 | 11.6 ($\pm 5.5$) | 882 ($\pm 97$) | 5.3 ($\pm 1.3$) | W1 |
| 29 | 1.2 ($\pm 0.8$) | 335 ($\pm 40$) | 1.7 ($\pm 0.7$) | W1 |
| 90 | 3.8 ($\pm 1.2$) | 648 ($\pm 80$) | 3.3 ($\pm 0.9$) | W1 |
| 141 | 0.9 ($\pm 0.1$) | 457 ($\pm 37$) | 2.3 ($\pm 0.7$) | W1 |
| 153 | 2.1 ($\pm 4.9$) | 401 ($\pm 53$) | 1.9 ($\pm 0.7$) | LL |
| 315 | 7.2 ($\pm 1.4$) | 620 ($\pm 66$) | 3.5 ($\pm 0.9$) | W1 |

**Table 2:**
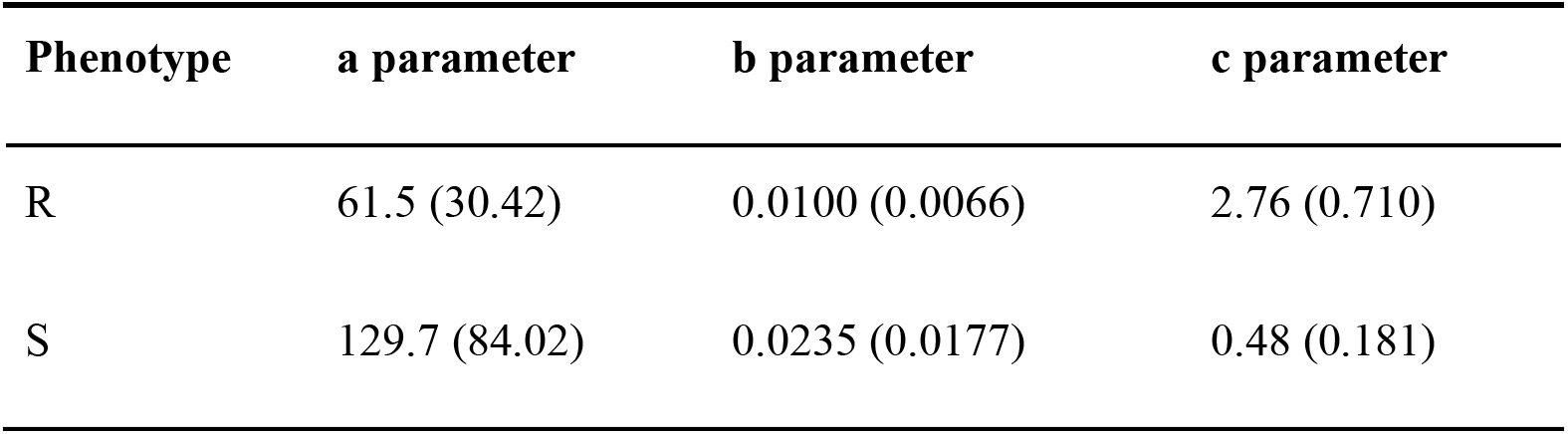
Hyperbolic model parameters for resistant (R) and susceptible (S) phenotypes. ***a*** indicates the maximum yield in the absence of competition; b indicates the competitive response of a phenotype to an increase in density and c denotes the relative competitive effect of the phenotype. Values in brackets are standard errors of the mean

### Gene amplification correlated with resistance to glyphosate

Higher ***EPSPS*** gene copy numbers were present in resistant seed families with mean values ranging between 3.8 – 11.6 copies. Moreover, there was a significant positive relationship between average seed family ***EPSPS*** relative gene copy number and seed family LD_50_ (Figure 4) (f-statistic= 22.99; df=1; 4, ***p*** =0.0087, R^2^ = 0.85). The linear model estimates that each gene copy increased glyphosate LD_50_ by 44.3 g ai ha^−1^. Increased ***EPSPS*** gene copy number was associated with enhanced glyphosate resistance in the ***A. tuberculatus*** Renville population. Target site sequencing of the ***EPSPS*** gene confirmed the absence of non-synonymous mutations in the active site of resistant individuals (data not shown). Furthermore, there was no significant impairment of translocation of ^14^C labelled glyphosate away from the site of application in resistant individuals (Supp. Figure 1).

**Figure 4:**
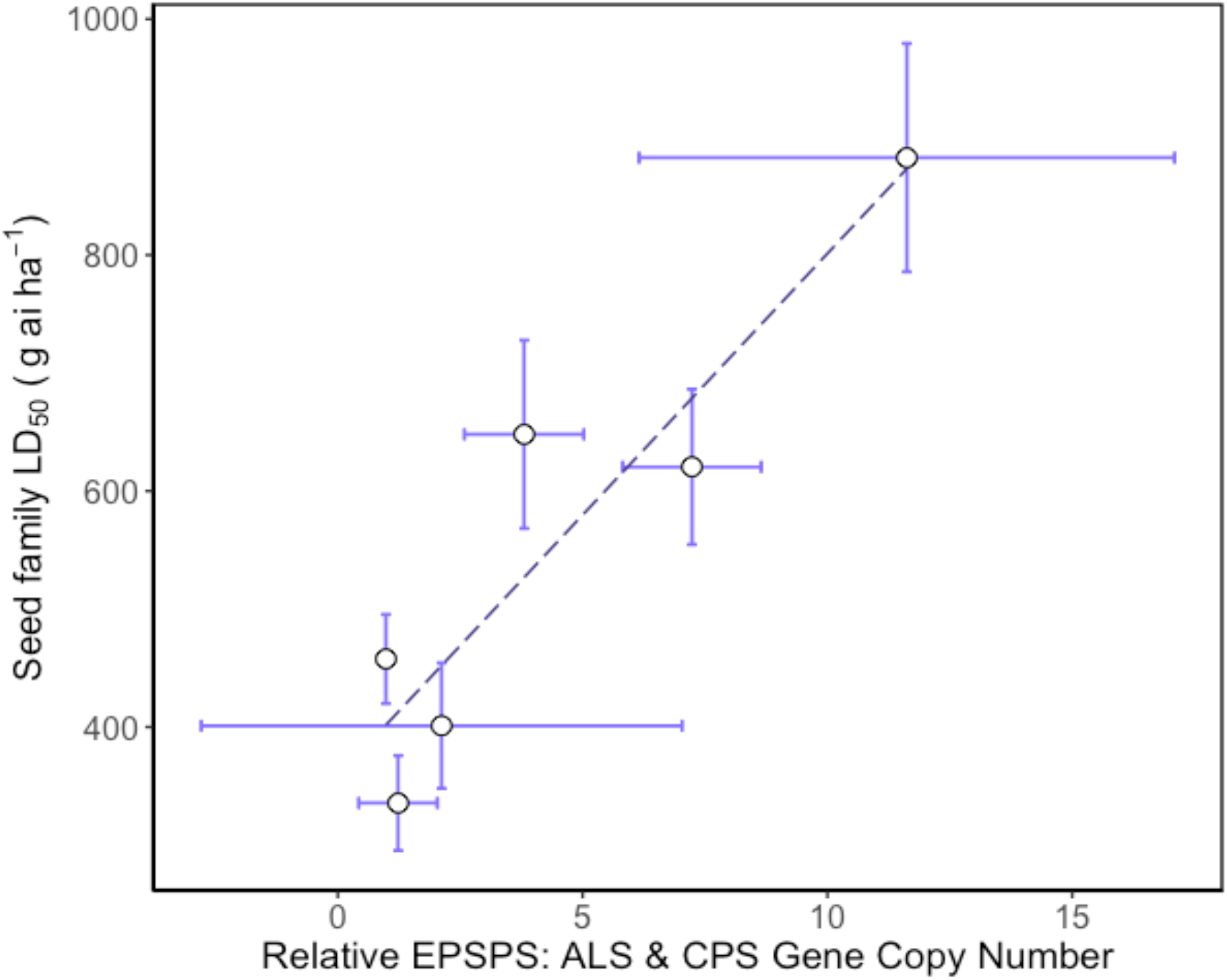
Relationship between relative mean *EPSPS* gene copy number and glyphosate resistance phenotype in *Amaranthus tuberculatus* seed families. Mean 5-enolpyruvylshikimate-3-phosphate synthase (***EPSPS***) gene copy number is relative to carbamoyl phosphate synthetase (***CPS***) *&* acetolactate synthase (***ALS***) reference genes. Error bars are standard errors of the mean.

### Decreased competitive effect of glyphosate resistant plants

Competitive effect and response were measured using a response surface experiment. No significant difference was observed between resistant (seed family 9) and susceptible (seed family 29) plant biomass in the absence of competition (parameter ***a***) nor in the response to competition (parameter ***b***). However, the competitive effect on neighbouring plants was greater for susceptible plants (parameter ***c***; *Z*_39_ = 3.107, ***p*** < 0.001). Resistant plants had a reduced competitive effect when compared to susceptible plants, where a substitution rate of 2.76 was determined for resistant phenotypes and 0.48 for susceptible phenotypes. Explicitly, 0.48 susceptible plants are required to have an equal competitive effect to that of a single resistant plant and 2.76 resistant plants are required to have an equal competitive effect to a single susceptible plant. The relative competition dynamics of resistant and susceptible phenotypes are shown in Figure 5 and Table 2.

**Figure 5:**
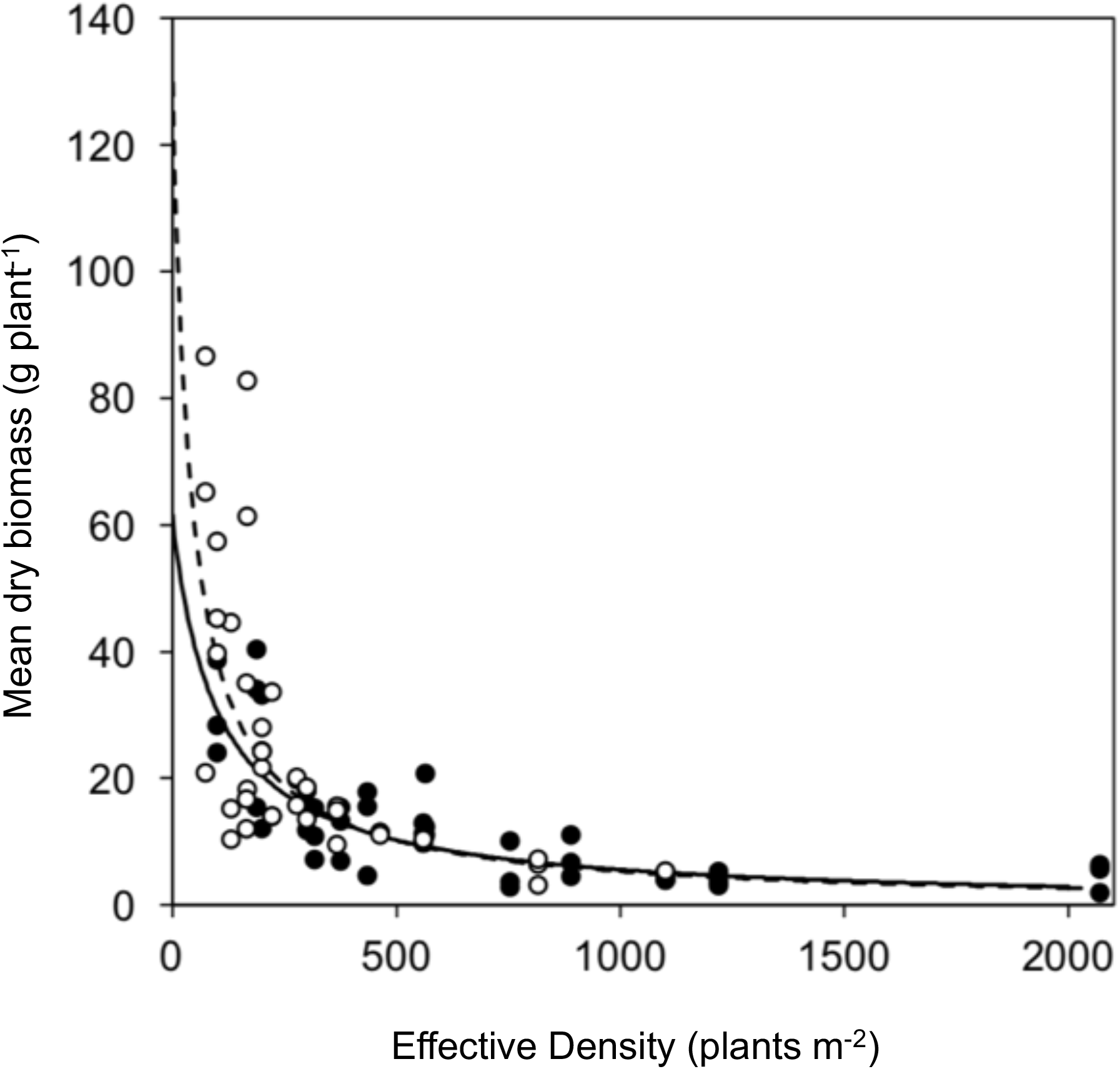
Vegetative growth of a glyphosate sensitive and resistant seed family in response surface competition experiment. Symbols represent mean biomass of resistant (black) and susceptible (white) phenotypes from replicate planting boxes at each effective density. Curves show fitted hyperbolic models for resistant (solid) and the susceptible (dashed)phenotypes.

### Interspecific competition influences resource allocation

Overall competition between the maize and ***A. tuberculatus*** seed families was assessed; Four ***A. tuberculatus*** plants reduced the above-ground biomass of one maize plant by 42 % (t78=-0.07, ***p***<0.001) whereas maize reduced the above ground biomass of ***A. tuberculatus*** by 37 % (t118 = −8.24, ***p*** <0.001). ***A. tuberculatus*** seed families with contrasting resistance levels were all observed to consistently divert resources away from vegetative growth in order to allow 10% more resource allocation towards reproduction in the presence of maize (t_118_ = −7.44, ***p*** <0.001). This resource allocation was observed across all seed families.

### Trade-off associated with resistance

A neighbourhood design competition experiment allowed the study of intraspecific and interspecific resistance costs. ***A. tuberculatus*** seed families were selected to represent a continuum of glyphosate resistance levels (and ***EPSPS*** gene copy number), thus making it possible to establish the extent of the trade-off between resistance benefit and resistance cost. Significant negative correlations were observed between above ground biomass in the absence of glyphosate application (a measure of resistance cost) and glyphosate LD_50_ (resistance benefit) (r=-0.87, ***p*** <0.05; Suppl. Figure 2) and gene copy number (GCN) (r=-0.83, ***p***<0.05; Figure 6) illustrating a significant quantitative resistance cost associated with enhanced GCN under intra-phenotypic competition. The relationship between reproductive biomass and LD_50_ followed the same trend (r= −0.87, ***p*** <0.05; Suppl. Figure 2), with a non-significant negative relationship observed between GCN and reproductive biomass (r=-0.76, ***p*** = 0.08; Figure 6). There was no relationship between seed family resistance level and the competitive effect of ***A. tuberculatus*** on maize and maize had an equal competitive effect on all ***A. tuberculatus*** seed families. Thus, the fitness cost established by these experiments was not evident under interspecific competition.

**Figure 6:**
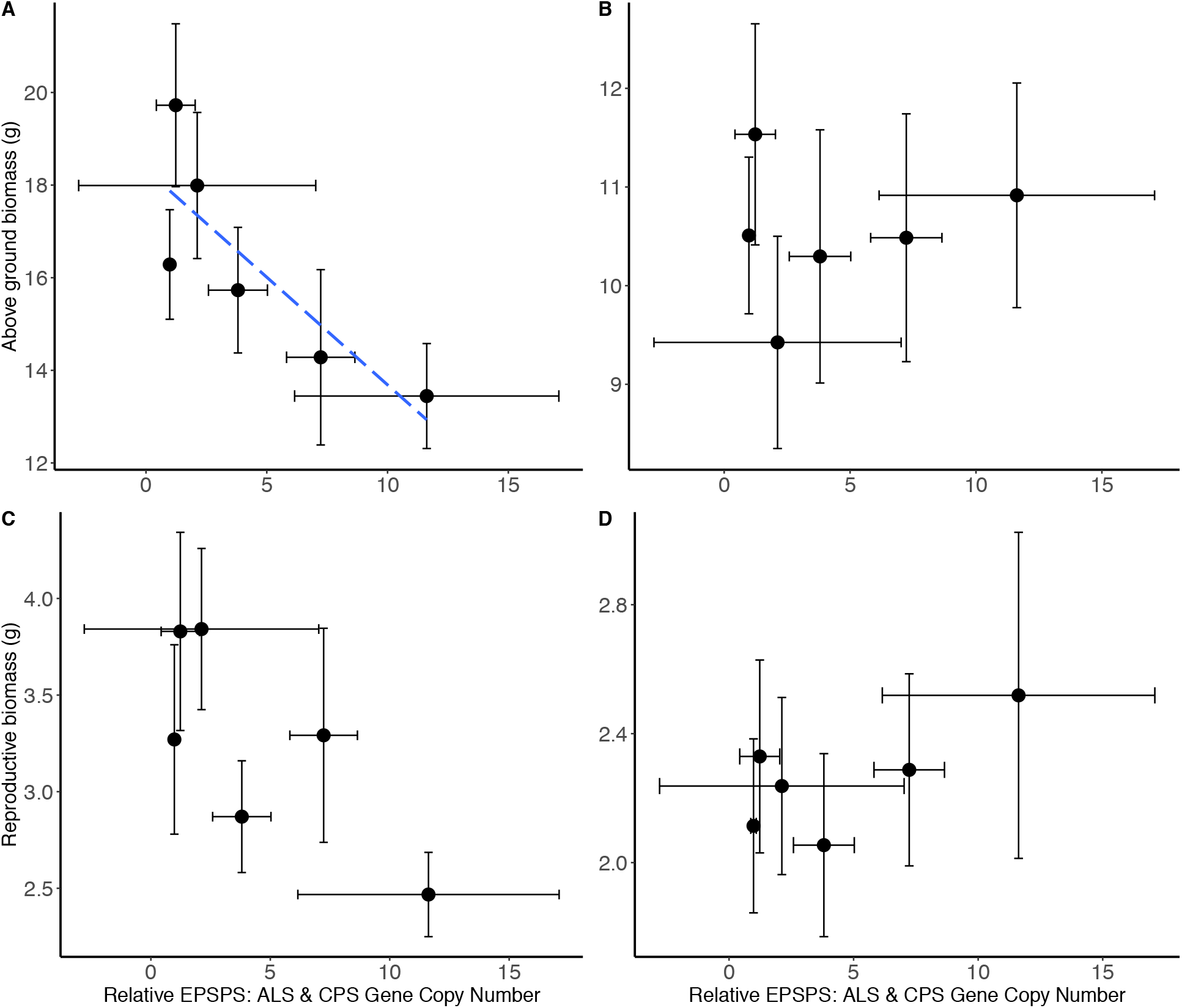
Biomass of common waterhemp seed families grown in neighbourhood design experiment. The relationship between ***Amaranthus tuberculatus*** seed family average above ground biomass (A & B) and reproductive biomass (C & D) compared to the average 5-enolpyruvylshikimate-3-phosphate synthase (***EPSPS***) gene copy number relative to carbamoyl phosphate synthetase (***CPS***) ***&*** acetolactate synthase (***ALS***) reference genes. A & C represent ***A. tuberculatus*** plant measures when grown under intra-phenotypic competition. the resistance level B & D represent ***A. tuberculatus*** measures when plants are grown in interspecific competition with maize. Error bars are standard errors of the mean.

## Discussion

The primary mechanism of glyphosate resistance in the common waterhemp population was gene amplification of the ***EPSPS*** target site. Seed families with contrasting glyphosate resistance status were successfully generated and higher ***EPSPS*** gene copy numbers (GCN) were associated with an increase in glyphosate resistance. Higher GCN were also associated with a growth penalty when grown under intra-phenotypic competition. This trade-off was mitigated in the presence of interspecific competition with maize. Further study of the seed families with extreme phenotypes revealed that the most resistant ***A. tuberculatus*** individuals had a lower competitive effect on neighbouring plants than the most susceptible seed family.

### Confirmation of *EPSPS* gene amplification based glyphosate resistance

***EPSPS*** gene amplification has been shown to be the main mechanism of glyphosate resistance found in ***A. tuberculatus*** populations (Murphy ***et al.*** 2019). ***EPSPS*** gene amplification has also been confirmed as the mechanism of evolved glyphosate resistance in ***Kochia scoparia***, ***Lolium multiflorum*** and ***Amaranthus spp.*** (Gaines et al. 2010, Salas et al. 2012, Nandula et al. 2014, Wiersma et al 2015). In ***A. tuberculatus EPSPS*** gene amplification has led to increased ***EPSPS*** transcripts and increased protein abundance, indicating that the additional gene copies are likely to be functional (Lorentz ***et al.*** 2014). No target site mutations nor conclusive evidence for the impaired translocation of glyphosate was observed in our study population. Nonetheless, target site mutations and reduced movement of glyphosate have both evolved in ***A. tuberculatus*** to result in glyphosate resistance (Nandula ***et al.*** 2013). ***EPSPS*** gene amplification is the main mechanism observed in our glyphosate resistant ***A. tuberculatus*** population and we found these copies led to an increase in resistance. The nature of this mechanism allowed the generation of a genotypic spectrum of seed families to assess the relationship between gene copy number and fitness.

The multiple ***EPSPS*** genes in common waterhemp have been physically mapped to discrete gene clusters on two homologous chromosomes within pericentromeric regions (Dillon ***et al.*** 2017). The localisation of these genes to pericentromeric regions indicates transposon-mediated local tandem gene duplication (Slotkin & Martienssen 2007; Tamaru 2010; Dillon ***et al.*** 2017; Gaines ***et al.*** 2019). By contrast, multiple gene copies of ***EPSPS*** in ***A. palmeri*** have arisen through extrachromosomal circular DNA (eccDNA) structures (Koo ***et al.*** 2018). Interspecies hybridisation between ***A. palmeri*** to ***A. tuberculatus*** is one potential avenue for the transmission of the gene amplification resistance mechanism (Dillon ***et al.*** 2017). Indeed hybridisation within the Amaranthus genus has resulted in such transmissions ***(Nandula et al. 2014)***(Oliveira 2017). However, all of the evidence to date confirms that the eccDNA mechanism of gene duplication in ***A. palmeri*** is independent from the tandem gene duplication observed in ***A. tuberculatus (Gaines, Patterson & Neve 2019)***. Thus, interspecies hybridisation has not transferred the mechanism between the two species, but instead convergent evolution has produced two different mechanisms of gene amplification to combat the same selection pressure. This finding highlights the importance of different mechanisms of rapid evolution within plants to overcome biotic and abiotic stresses (Patterson ***et al.*** 2018).

### Resistance individuals exhibit a reduced competitive effect

The comparison of inter-phenotypic and intra-phenotypic competition between resistant and susceptible seed families was achieved through a response surface experiment. Susceptible ***A. tuberculatus*** individuals exhibited a greater competitive effect than resistant individuals, manifest as increased competitive suppression of neighbouring plants across different densities and proportions. A greater competitive effect indicates that resources are allocated to interference competition, or rather the inhibition of a neighbours ability to acquire resources. Our results illustrate that the ability to suppress the capture of local resources by neighbouring plants is an effective competitive strategy under intraspecific competition. These findings support the results observed in the neighbourhood design experiments. Similar studies investigating herbicide resistant ***Lolium rigidum*** have found no differences (Pedersen ***et al.*** 2007) and extensive costs (Vila-Aiub ***et al.***, 2009a) in competitive parameters.

### Intraspecific competition found to produce a resistance penalty

This work showed a growth and fitness trade-off across a glyphosate resistance spectrum of common waterhemp seed families under intra-phenotypic competition. Similarly. seed families with higher GCN were associated with a growth penalty under intra-phenotypic competition. Our results complement studies which employ alternative methods to elucidate putative fitness costs. Specifically, populations of ***A. tuberculatus*** were grown under fallow-like settings in the absence of selection and genotype frequencies were observed over six generations. Under these conditions a 10-fold reduction was observed in the number of individuals harbouring multiple ***EPSPS*** gene copies (Wu, Davis & Tranel 2018). By contrast, the ***EPSPS*** target site mutation, which also endowed glyphosate resistance, was found to persist where gene amplification declined (Wu ***et al.*** 2018). Thus, it appears that the ***EPSPS*** gene amplification mechanism is associated with a penalty in common waterhemp. Interestingly, no trade-off was associated with amplified ***EPSPS*** gene copy numbers in ***A. palmeri*** plants grown at two different densities (Vila-Aiub ***et al.*** 2014). However, glyphosate resistant palmer amaranth populations were seen to have a lower competitive ability across four crops (Chandi ***et al.*** 2013). As mentioned above ***A. palmeri*** contains an eccDNA mediated mechanism of gene amplification rather than tandem gene duplication. It is clear that the presence of an ***EPSPS*** gene amplification resistance cost may be influenced by weed species and the genomic mechanism that has mediated an increase in gene copy numbers.

### Interspecific competition was able to mitigate the intraspecific resistance penalty

The resistance penalty we observed under conditions of intra-phenotypic competition was not enhanced but mitigated in the presence of interspecific competition with maize. This finding indicates that, in spite of the penalties observed in intraspecific competition, the factors influencing the frequency of genotypes containing gene amplification mediated resistance are complex. These artificial experiments suggest that the expression of the trade-off is influenced by genotype x environment interactions. In fact, it is not the first time that such a phenomenon has been observed, where the presence of a crop influences the expression of a cost. When glyphosate resistant ***Lolium rigidum*** was grown in isolation or under low wheat competition, a fitness cost was observed resulting in a 7.5 % reduction in seed production. However, under high crop densities, the fitness cost was mitigated (Pedersen ***et al.*** 2007). Subsequent studies showed that the cost observed under low densities was maintained in the field under the absence of selection, revealing a 34% reduction in the resistant phenotype over 3 years (Preston & Wakelin 2008). The dominance and suppressor model of competition states that size differences are exacerbated under competition (Harper 1977; Weiner 1985). Here a larger plant, or one with a faster growth rate, was able to seize the majority of resources resulting in a greater ability to suppress the growth of competitors (Weiner 1985). We hypothesise that the variance in the exponential growth rates between maize and common waterhemp contributed to asymmetrical capture of resources resulting in size hierarchies (Schwinning & Weiner 1998). Such hierarchies have been found to result in increased size inequality at higher densities (Weiner 1985). Indeed the development of size hierarchies at high plant densities can be explained by unequal light interception (Schwinning & Weiner 1998) and also differences in growth rate (Schmitt, Ehrhardt & Cheo 1986). It is possible that hierarchies may have resulted in partial size asymmetry whereby the maize received a disproportionate share of the resources (Schwinning & Weiner 1998) thus uniformly restricting the growth of ***A. tuberculatu***s. Furthermore, the dominance / suppressor model comes into play at low light levels (Schmitt ***et al.*** 1986), and the polytunnel growth environment did not supply additional light, thus potentially assisting maize to act as the dominant competitive suppressor species.

### Exploitation of fitness penalties

Fitness cost information can be used in a more direct way to predict the efficacy of management practices in reducing the level of resistant genotypic frequencies observed within the field. This can be achieved through the accentuation of fitness costs which are found to be associated with herbicide resistance (Cousens & Mortimer 1995; Jordan 1999). It is evident that the trade-offs in life history traits associated with herbicide resistance may be selected against under specified management conditions leading to a reduction in the frequency of resistant phenotypes (Vila-Aiub ***et al.*** 2005a). Mathematical modelling to predict the evolution and spread of resistance can be used to tailor management practices and mitigate the risk of an herbicide resistance epidemic (Vila-Aiub ***et al.*** 2005b). Fitness benefit and cost parameters are required to model the dynamics of resistance evolution under specified management strategies. Modelling the impact of different management strategies on the evolution of glyphosate resistance in ***Amaranthus spp.*** has been used to predict resistant genotype frequency overtime (Neve ***et al.*** 2011; Liu ***et al.*** 2020). The adoption of the low resistance risk management strategies can be used to mitigate the risk of glyphosate resistance evolution (Neve ***et al.*** 2011; Liu ***et al.*** 2020). These two latter Amaranth models do not account for fitness costs. However, other weed models do cater for fitness costs associated with herbicide resistance (Colbach ***et al.*** 2016) where the addition of pleiotropic parameters enhanced the accuracy of the model predictions.

## Conclusion

Glyphosate resistant ***A. tuberculatus*** plants showed a reduced competitive ability to capture resources resulting in a negative trade-off between resistance and fitness in the absence of selection, however this resistance penalty was seen to be mitigated in the presence of maize, a dominant suppressor species. Ultimately, the frequency of genotypes containing gene amplification mediated glyphosate resistance is unlikely to decline in the absence of selection, within a maize production system.

## List of Abbreviations

ALS: acetolactate synthase
CPS: carbamoyl phosphate synthetase
Δ CT: Change in cycle threshold
eccDNA: extrachromosomal circular DNA
*EPSPS*: 5-enolpyruvylshikimate-3-phosphate synthase
LD_50_: Lethal dose 50
R: Resistant
S: Susceptible
TE: transposable element
qPCR: quantitative polymerase chain reaction

## Acknowledgements

This paper is based on work conducted as part of a PhD project funded through the Biotechnology and Biological Science Research Council and Syngenta. Confirmatory qPCR was performed at Rothamsted Research.

## Author Contributions

Conception and design of project – PN, SSK, HC

Writing manuscript – HC

Conducting phenotypic screens, dose response experiments, translocation assessment, competition experiments and statistical analysis – HC

Dose response advice - SJH

Radio labelling tutoring - AH

DNA extraction and Q-PCR – LN HC RPD

## Supplementary Methods

### Target site sequencing

Sanger sequencing was used to determine potential ***EPSPS*** target site mutations for 5 resistant and 3 susceptible parental plants. PCR reactions were Ready-to-go Taq beads (Amersham Biosciences), 22 μl water, 1 μl of each primer (10 mM), 1 μl template. Primer sequences in Supp. Table 1. PCR conditions were 95 oC for 5 min, 45 cycles of 95 oC for 15 sec, 60 oC for 1 min and 72 oC for 1 min. Then 72 oC for 1 min. Sequences were generated through sequencing.

### Impaired glyphosate translocation

^14^C glyphosate was applied using a Hamilton syringe in 20 x 0.2 μl droplets to resistant and susceptible plants pre-treated with 840 g ai ha^−1^ glyphosate. An average of 5517 Bq ^14^C was applied to each plant. Twelve replicate resistant and susceptible plants were assessed across the eight treatments; 0, 6, 24 and 72 hours post treatment. Plants were dissected based on tissue type (application site, above treated area, meristem, remaining foliage, stem, and roots) and combusted in a Zinsser Robox 192 Biological oxidizer (OX500; Harvey). Evolved ^14^CO_2_ was trapped in Oxysolve C400 and released through scintillation fluid to allow quantification. Translocation was calculated as the proportion of glyphosate transported away from the site of application as a percentage of the residue recovered at time point zero.

**Supplementary Figure 1.**
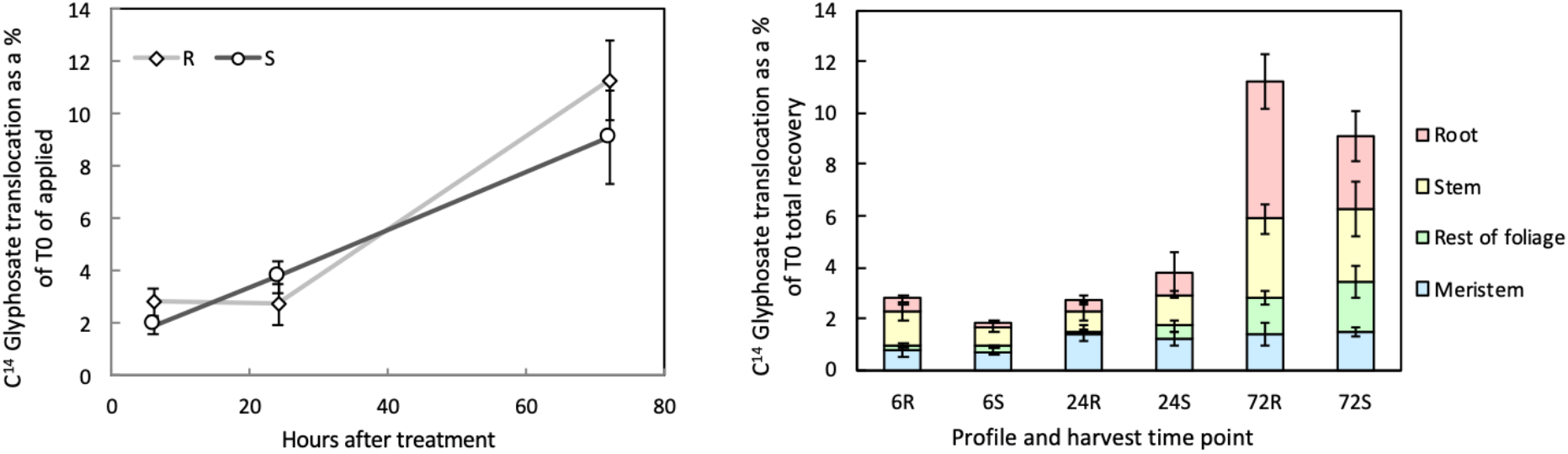
Glyphosate resistance mechanism of resistance in common waterhemp. a) ^14^C labelled glyphosate translocated away from the site of application in the resistant (R) and susceptible (S) ***A. tuberculatus*** plants relative to the ^14^C glyphosate recovery at time point zero (T0) b) The proportions of ^14^C labelled glyphosate translocated away from the site of application to the root, stem, rest of the foliage and meristem, relative to the ^14^C glyphosate recovery at time point zero (T0)

**Supplementary Figure 2.**
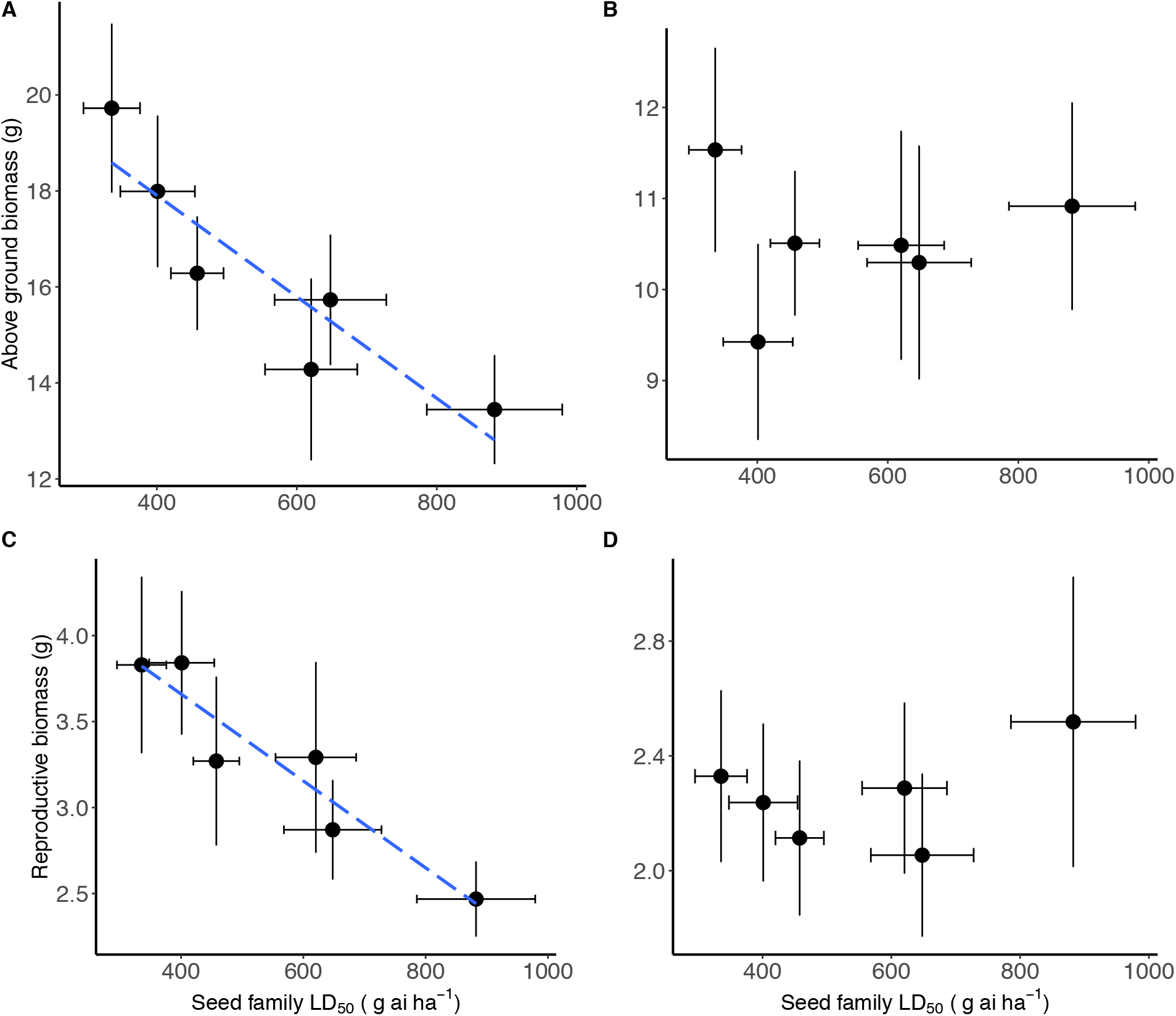
Biomass of common waterhemp seed families grown in neighbourhood design experiment. The relationship between ***Amaranthus tuberculatus*** seed family average above ground biomass (A & B) and reproductive biomass (C & D) compared to the resistance level glyphosate lethal dose required to kill 50 % of individuals (LD_50_). A & C represent ***A. tuberculatus*** plant measures when grown under intra-phenotypic competition. the resistance level B & D represent ***A. tuberculatus*** measures when plants are grown in interspecific competition with maize. Error bars are standard errors of the mean.

**Supplementary Table 1.**
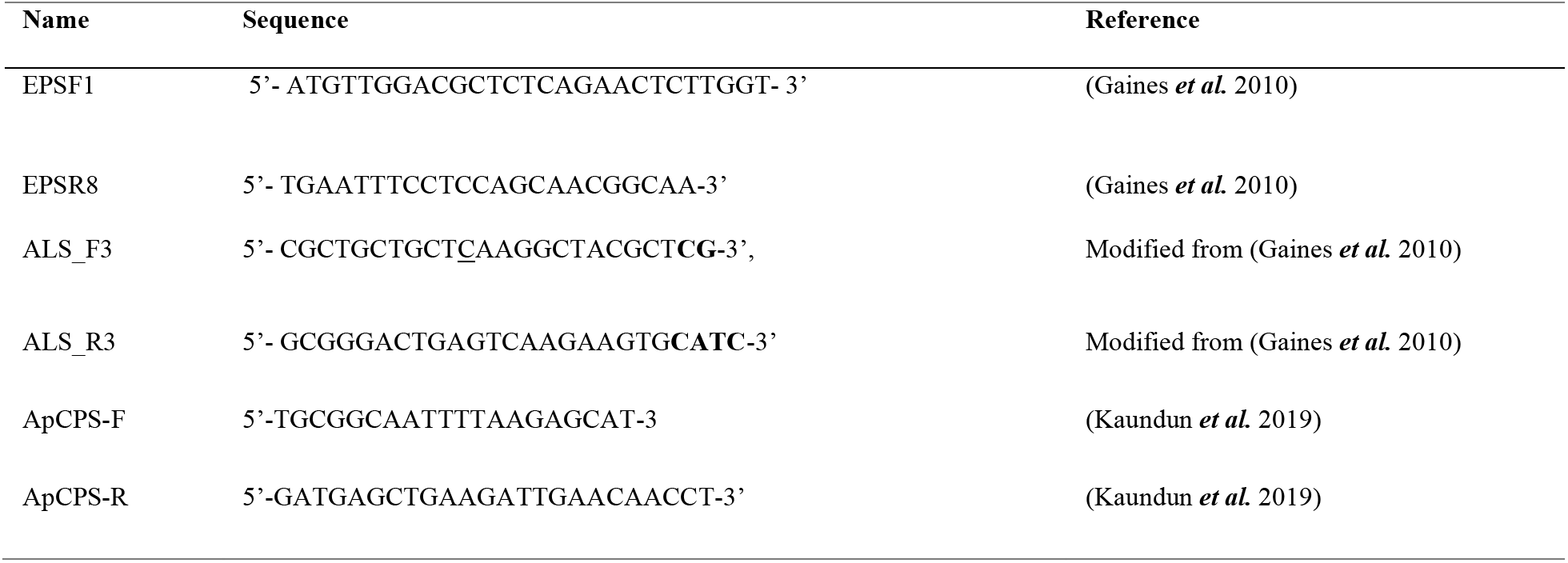
Primer sequences for qPCR.

## Notes

### Competing Interest Statement

SSK, RPD, AH and SHJ were employed by company Syngenta
The remaining authors declare that the research was conducted in the absence of any commercial or financial relationships that could be construed as a potential conflict of interest.

